# Optimizing the Accuracy of Cortical Volumetric Analysis in Traumatic Brain Injury

**DOI:** 10.1101/822148

**Authors:** Bram R. Diamond, Christine L. Mac Donald, Samuel B. Snider, Bruce Fischl, Kristen Dams-O’Connor, Brian L. Edlow

**Author notes:** co-senior authors (Kristen Dams-O’Connor, Brian L. Edlow). Correspondence to: Kristen Dams-O’Connor, Ph.D. **Author Contact Information**: Bram R. Diamond, address: Athinoula A. Martinos Center for Biomedical Imaging, 149 Thirteenth Street, Charlestown, MA 02129. Christine L. Mac Donald, address: University of Washington School of Medicine, Department of Neurological Surgery, 325 9^th^ Ave, Box 359924, Seattle, WA 98104. Samuel B. Snider, address: Massachusetts General Hospital, 175 Cambridge St – Suite 300, Boston, MA 02114. Bruce Fischl, address: Athinoula A. Martinos Center for Biomedical Imaging, 149 13^th^ Street, Charlestown, MA 02129. Kristen Dams-O’Connor, address: Brain Injury Research Center, Department of Rehabilitation Medicine, Department of Neurology, Box 1163, One Gustave L. Levy Place, New York, NY 10029. Brian L. Edlow, address: Massachusetts General Hospital, 175 Cambridge St – Suite 300, Boston, MA 02114.

## Abstract

Cortical volumetric analysis is widely used to study the anatomic basis of neurological deficits in patients with traumatic brain injury (TBI). However, patients with TBI-related lesions are often excluded from analysis, because cortical lesions may compromise the accuracy of reconstructed surfaces upon which volumetric measurements are based. Here, we propose a novel FreeSurfer-based lesion correction method and illustrate its impact on cortical volume measures in patients with chronic moderate-to-severe TBI. We performed MRI in 87 patients at mean+/−SD 10.9+/−9.1 years post-injury using a T1-weighted multi-echo MPRAGE sequence at 1 mm resolution. Following surface reconstruction, we parcellated the cerebral cortex into seven functional networks using FreeSurfer’s standard pipeline. Next, we manually labeled vertices on the cortical surface where lesions caused inaccuracies and removed them from network-based cortical volumetric measures. After performing this lesion correction procedure, we measured the surface area of lesion overlap with each network and the percent volume of each network affected by lesions. We identified 120 lesions that caused inaccuracies in the cortical surface in 46 patients. In these 46 patients, the most commonly lesioned networks were the limbic and default mode networks (95.7% each), followed by the executive control (78.3%), and salience (71.7%) networks. The limbic network had the largest average surface area of lesion overlap (4.4+/−3.7%) and the largest percent volume affected by lesions (12.7+/−9.7%). The lesion correction method has the potential to improve the accuracy of cortical volumetric measurements and permit inclusion of patients with lesioned brains in quantitative analyses, providing new opportunities to elucidate network-based mechanisms of neurological deficits in patients with TBI.

## Introduction

Cortical volumetric analysis with FreeSurfer^1, 2^ is widely used to study the neuroanatomic basis of cognitive, behavioral, and motor deficits in patients with traumatic brain injury (TBI).^3–6^ However, cortical lesions caused by TBI pose major challenges to FreeSurfer’s standard automated magnetic resonance imaging (MRI) processing pipeline. Lesions often compromise the accuracy of the cortical surfaces that are reconstructed and used by FreeSurfer to generate volumetric measurements.^4, 6, 7^ As a result, TBI imaging studies have historically excluded patients with large focal lesions.^5, 8^ Development of a tool that accounts for lesions in cortical volumetric analysis is needed to prevent the systematic exclusion of patients with large cortical lesions and to ensure that TBI imaging studies are generalizable across the full spectrum of cortical pathology. Moreover, integration of such a tool into the FreeSurfer software platform would create new opportunities to study network-based mechanisms of disease^9, 10^ using canonical atlases.^11^

Here, we propose a novel FreeSurfer-based lesion correction method and illustrate its impact on cortical volumetric measures in patients with chronic TBI. The lesion correction method differs in several ways from the standard FreeSurfer approach to editing reconstructed cortical surfaces. Standard cortical segmentation using FreeSurfer relies on the assumption that the brain has normal anatomy and that any surface inaccuracies are related to the FreeSurfer processing pipeline. However, in patients with cortical lesions caused by TBI, FreeSurfer’s reconstruction of the cortical surface can be grossly inaccurate due to focal encephalomalacia and distorted anatomy. This methodological limitation of the standard FreeSurfer editing approach is the main motivation for the lesion correction method proposed here. The new method makes no assumptions about lesioned cortical surface anatomy, and it minimizes bias by requiring the manual rater simply to identify inaccuracies without changing the surfaces. In this study, we use the lesion correction method to assess the topology of lesion overlap with functional brain networks and to characterize inter-network differences in lesion burden. We also distribute the lesion correction method to the academic community to facilitate future studies of network-based mechanisms of neurological deficits in patients with TBI.

## Methods

### Patients

Between May, 2014 and January, 2019, we prospectively enrolled 141 patients with a history of TBI at two academic medical centers as part of the Late Effects of TBI (LETBI) study.^12^ Patients were included if they had sustained a moderate-to-severe TBI at least one year prior to enrollment. We characterized TBI severity based on the United States Department of Defense classification system,^13^ as detailed in the Supplementary Material. Of the 141 enrolled participants, 98 completed an MRI scan (see CONSORT diagram in Supplementary Fig. S1).

### MRI data acquisition

Patients at Mount Sinai were scanned using a Siemens Skyra (Siemens Medical Solutions, Erlangen, Germany) 3 Tesla (T) MRI scanner with a 32-channel head coil for signal reception, and patients at University of Washington were scanned using a Philips Achieva 3T MRI scanner with a 32-channel head coil.^1^ Patients underwent standardized MRI using a T1-weighted multi-echo MPRAGE (MEMPRAGE)^14^ sequence with 1mm isotropic voxels. All LETBI sequences were designed to maximize consistency with the National Institutes of Health Common Data Elements for TBI Neuroimaging.^15^

### MRI processing

We first processed all MEMPRAGE data using the standard FreeSurfer pipeline (version 6.0) for cortical surface reconstruction and cortical volume estimation.^1^ We used the “big ventricles” function to optimize automatic segmentation for a patient population with enlarged ventricles. In accordance with FreeSurfer recommended best practices (https://surfer.nmr.mgh.harvard.edu/fswiki/FsTutorial/TroubleshootingDataV6.0), we visually inspected output files, made manual edits to the white matter segmentation, and added control points. To ensure that the lesion correction method would be tested in an unbiased manner, we did not manually edit regions bordering cortical lesions. We then resampled the Yeo 7-Network resting-state functional connectivity atlas^11^ onto each patient’s reconstructed cortical surface using FreeSurfer’s surface-based registration tool (https://surfer.nmr.mgh.harvard.edu/fswiki/mri_surf2surf).^16^

### Quality Assessments

We performed visual quality assessment for all 98 scans based upon delineation of grey-white matter boundaries and the accuracy of the FreeSurfer-generated surfaces. We defined scan quality using an integer scale: 0 = scan excluded because FreeSurfer failed to complete the processing pipeline; 1 = scan excluded because surface inaccuracies would have required major manual edits; 2 = scan included because only minor manual edits required; 3 = scan included without requiring manual edits. For any scan that received a score of 1 by the primary rater (B.R.D.), a second rater (B.L.E.) reviewed the scan to achieve consensus. Our primary method for determining scan inclusion was qualitative visual assessment because inaccurate FreeSurfer-based segmentations can confound quantitative measurements. Nevertheless, we performed quantitative assessments of signal-to-noise ratio (SNR) and contrast-to-noise ratio (CNR) and tested for correlations with visual assessments of scan quality, as detailed in the Supplementary Materials.

### Lesion identification and classification

We next assessed each MRI scan for focal lesions causing encephalomalacia of the cerebral cortex (Fig. 1, top row).^15^ All such lesions were considered for subsequent lesion correction analysis and classified according to the cortical network(s) with which they overlapped. To ensure robust and reproducible methods for lesion identification, we performed an inter-rater reliability analysis among three investigators who identified lesions in a randomly selected group of 20 MRI scans and calculated lesion volumes using the standard ABC/2 method.^17^ Two investigators were board-certified neurologists with fellowship training in Neurocritical Care (B.L.E. and S.B.S.) and one was a research technician (B.R.D.).

**Figure 1.**
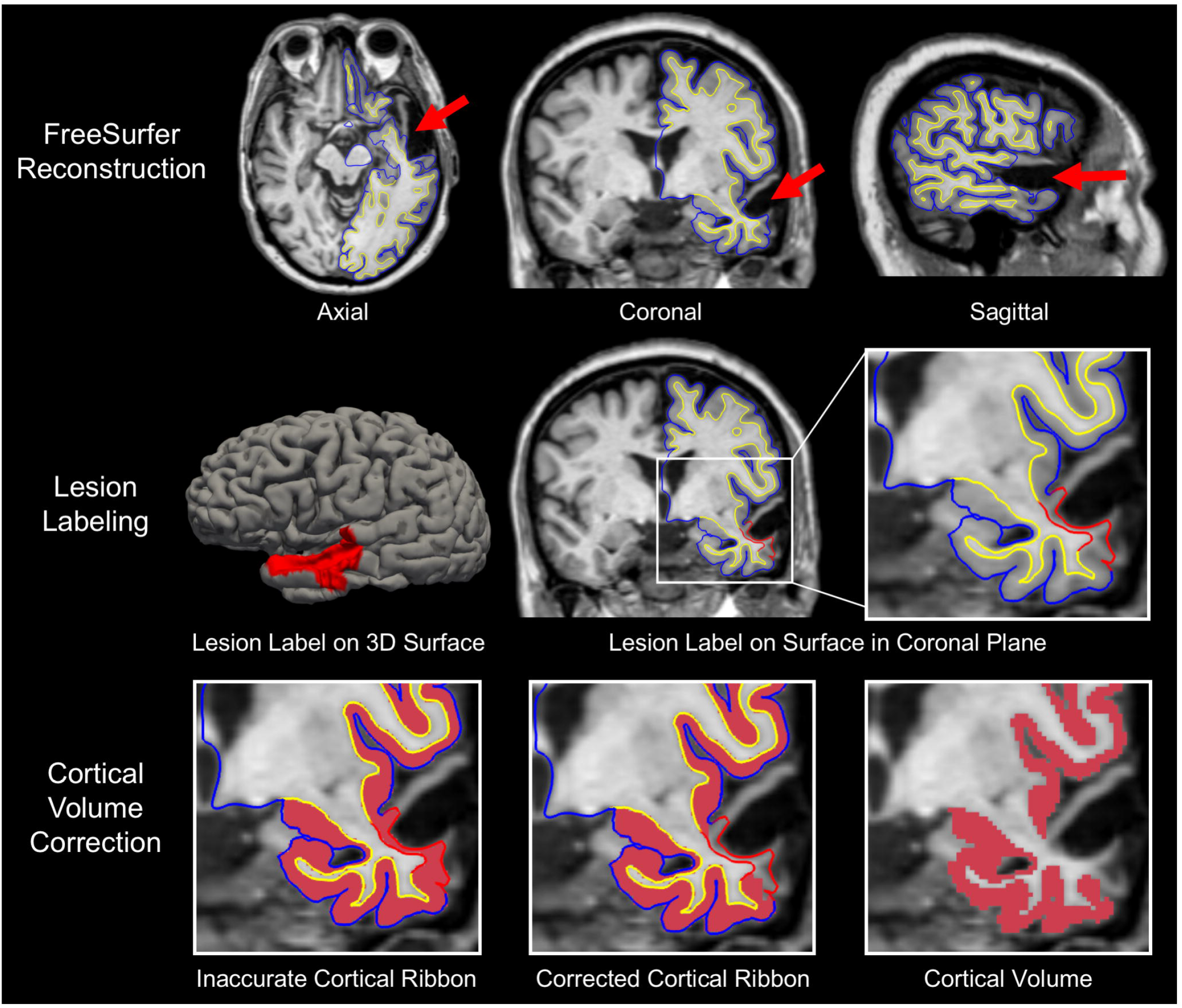
Overview of Lesion Correction Method. Row 1: Axial, coronal, and sagittal T1-weighted images of a representative patient with traumatic brain injury. FreeSurfer reconstructions of the cortical surface (blue line) and grey-white surface (yellow line) are used to visually identify regions where a cortical lesion (red arrows) caused surface inaccuracies. Row 2: We manually outlined lesions by labeling inaccurate vertices on the cortical surface (left image). This surface inaccuracy (labeled in red) is shown in the coronal plane in the middle image and the right, zoomed image. The red label passes through lesioned, encephalomalacic tissue. Row 3: To correct for the inaccuracy in the surface label at the site of the lesion, we remove the volume of cortex within the lesion label and perform cortical volumetric measures that exclude the lesioned tissue.

### Implementation of the lesion correction procedure

A detailed description of the methodological principles of the lesion correction procedure is provided in the Supplementary Material. To implement the procedure, we visually identified sites where FreeSurfer’s modeled surface mesh erroneously passed through subcortical tissue (Fig. 1, top row). Next, we manually labeled these surface-points to produce lesion-induced inaccuracy labels (Fig. 1, middle row). Finally, we applied these labels as exclusion masks to remove affected surface regions and calculate corrected cortical volumes (Fig. 1, bottom row).

After performing this lesion correction procedure, we used standard FreeSurfer tools to measure the average surface area overlap of lesion-induced inaccuracies with each network of the Yeo 7-Network atlas^11^ and the average percent volume change of each network caused by the lesion correction procedure (Fig. 2). There was no need to correct cortical volume measurements by total intracranial volume in this study because all network-based measures (i.e. % change in volume) were calculated at the single-subject level.

**Figure 2.**
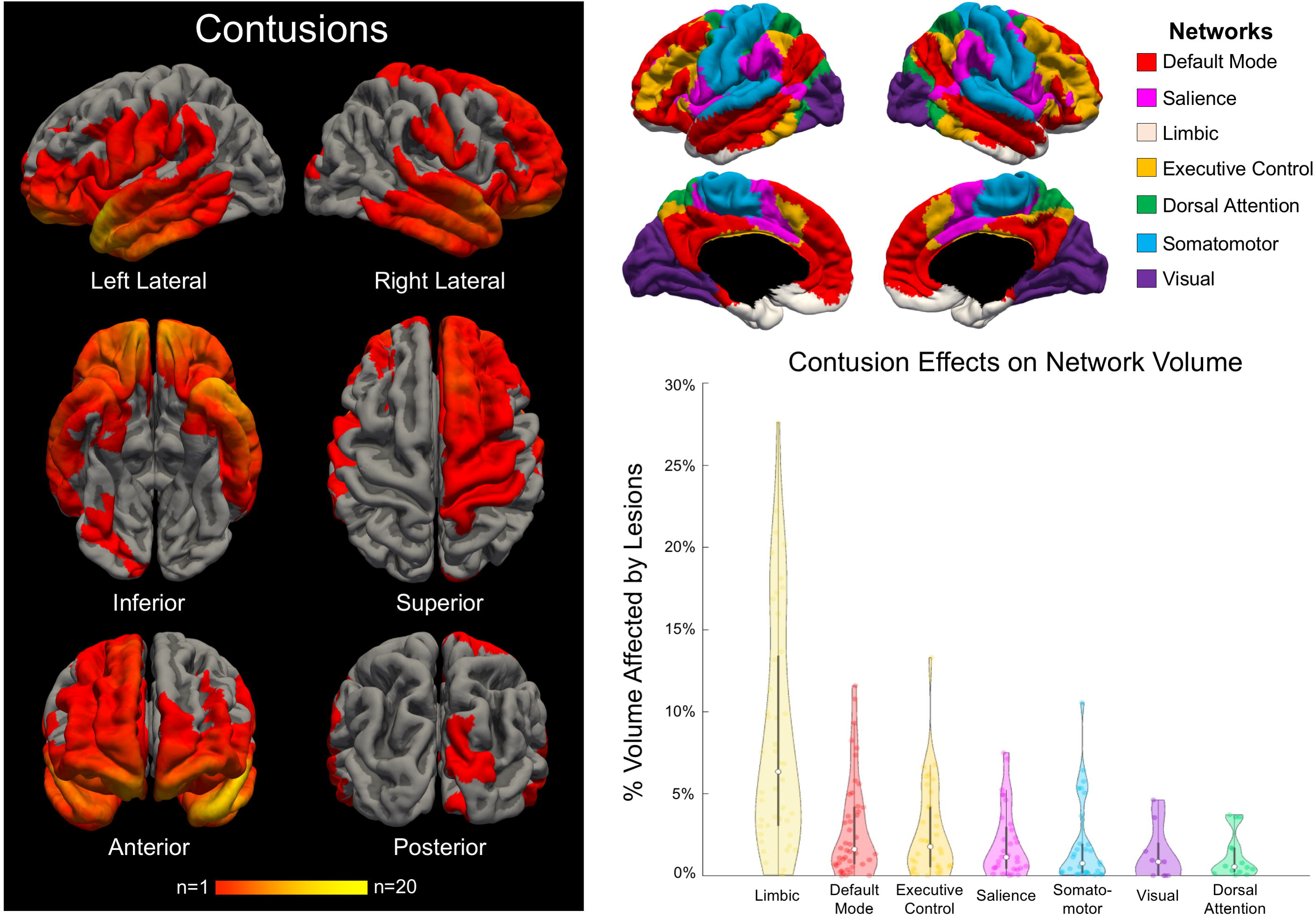
Lesion Topology and Network-based Lesion Effects on Cortical Volume. In the left panel, we show a heat map of cortical lesions for all 46 patients who had at least one lesion. The anatomic regions most commonly affected by cortical lesions were the frontal and temporal lobes, particularly the frontal poles, temporal poles and orbitofrontal regions. In the top right panel, we show the 7 functional networks from the Yeo atlas^11^ that were used to investigate network-specific lesion effects. In the bottom right panel, we show a violin plot demonstrating the changes in average cortical volume for each network after applying the lesion correction method. Lesion effects on average cortical volume varied between networks, with the limbic network showing the largest magnitude of decline in average cortical volume after application of the lesion correction method.

An overview of the lesion correction procedure is shown in Video 1, and additional methodological details are provided in the Supplementary Material. We also release all code used in the lesion correction procedure on https://github.com/ComaRecoveryLab/Lesion_Correction.

### Statistical analysis

We used the intraclass correlation coefficient to test interrater reliability for lesion volume measurements. We report descriptive statistics for the average percent cortical surface area and the average percent cortical volume affected by lesions for each network.

## Results

### Patient demographics and clinical characteristics

Due to the presence of severe anatomic distortions, two of the 98 patients’ scans did not complete FreeSurfer’s standard processing pipeline (visual assessment scores=0). Of the remaining 96 scans, nine received a visual assessment score of 1 by the two raters and were excluded, yielding a final sample size of 87 patients. The 87-patient cohort was comprised of 60.9% men, with a mean +/− SD age of 56.7 +/− 12.0 years. Injury severity was classified as mild (n=3), moderate (n=42), and severe (n=32); in 10 participants duration of LOC was unknown and records were not available. The duration from most recent TBI to MRI was 10.9 +/− 9.1 years. Additional clinical and demographic data, as well as SNR and CNR data, are provided in Supplementary Tables 1 and 2.

### Interrater Reliability

The intraclass coefficient between the two physician raters across 20 datasets was 0.99 [95% Confidence Interval 0.98, 0.99]. The intraclass coefficients between the physician raters and the technician rater for these same datasets were 0.95 [0.91, 0.97] and 0.96 [0.93, 0.98], respectively. Because sufficient inter-rater reliability was established in this test set (n=20; intraclass correlation coefficient > 0.9), all subsequent lesion identification was performed by the technician rater, B.R.D.

### Lesion characteristics and anatomic distribution

Forty-six of the 87 patients had at least one lesion that affected the accuracy of the FreeSurfer-modeled cortical surface. There were 120 total lesions, with a median of 2 lesions per patient (range 1 to 10). On average, lesions overlapped with 4.6 +/− 1.6 of the 7 networks. A group-level lesion topology map demonstrated an orbitofrontal and anterior temporal predominance of the lesions (Fig. 2, Videos 2 and 3).

### Network-based cortical surface area measures

The limbic and default mode networks were lesioned in the largest proportion of patients (44/46 scans, 95.7% incidence for both networks), followed by the executive control (78.3%), and salience (71.7%) networks. This large limbic lesion burden was observed despite the limbic network having the smallest average surface area of the seven functional networks across all patients (Supplementary Fig. S2). The largest mean percentage of lesion-network surface area overlap occurred within the limbic network (4.4 +/− 3.7% of total network surface area; Supplementary Table 3).

### Network-based cortical volume measures

When considering networks impacted by the lesion correction method in the 46 patients with cortical lesions, we observed a median decrease in network-based cortical volume of 3.4% (range <1.0% to 47.0%). The limbic network had the largest lesion-induced mean +/− SD percentage decrease in cortical volume (12.7 +/− 9.7%; Supplementary Table 4).

## Discussion

We introduce a new FreeSurfer-based method for cortical volumetric analysis in patients with lesions caused by TBI. We apply this method in a cohort of 87 patients with chronic moderate-to-severe TBI and show that lesion-induced cortical inaccuracies are not equally distributed within the brain’s functional networks. Rather, inaccuracies preferentially affected the limbic network, an observation consistent with prior pathology^18, 19^ and MRI^20^ studies showing that traumatic contusions commonly affect the orbitofrontal and temporal nodes of the limbic network. Implementation of the proposed lesion correction method will prevent the systematic exclusion of patients with cortical lesions from MRI volumetric studies and improve the generalizability of MRI studies across the full spectrum of cortical pathology.

These findings demonstrate the potential utility of the new lesion correction method for studying network-based mechanisms of cognitive, behavioral, and motor deficits in patients with TBI. For example, lesion-induced cortical volume changes within the limbic, default mode, and frontoparietal networks (the three most frequently lesioned networks) can be tested for correlations with symptoms that are putatively attributable to their dysfunction, such as behavioral dysregulation, altered self-awareness, and executive dysfunction, respectively. From a phenomenological standpoint, the application of the new lesion correction tool to large clinical-radiological-pathological databases being acquired by the LETBI,^12^ Transforming Research and Clinical Knowledge in TBI (TRACK-TBI),^21^ Collaborative European NeuroTrauma Effectiveness Research in Traumatic Brain Injury (CENTER-TBI),^22^ and other studies, has potential to elucidate pathological signatures of TBI phenotypic classification, with implications for clinical trial selection^23^ and prognostication.^10^

Several limitations should be considered when interpreting the results of this study. The lesion correction method relies upon an assumption whose validity is difficult to test: we assume that at sites of tissue distortion and encephalomalacia, the cortex is non-functional and therefore should be masked, or removed, from subsequent cortical volume measurements. This assumption is made with the recognition that definitive determination of the functional status of lesioned cortex is not possible solely with T1-weighted MEMPRAGE data. Nevertheless, the assumption that lesioned cortex is non-functional in the population studied here is strongly supported by visual inspection of the data, which reveals complete or near complete absence of cerebral cortex, as shown in Figure 1. In future multimodal experiments, the lesion correction method can be refined by analyzing the functional properties of lesioned cortex (e.g. with functional MRI or EEG). In future work, it may also be possible to integrate the lesion correction method with software programs that offer automated lesion detection, such as the ABC module extension of 3D Slicer.^24^ Moreover, the method can be used to measure point-wise and region-wise estimates of cortical thickness in unlesioned cortex by masking inaccurate regions of cortex.

## Conclusions

We demonstrate the impact of a new FreeSurfer-based lesion correction tool on cortical volumetric measures in 7 atlas-based functional networks, and we distribute this lesion correction tool to the academic community. We show that cortical lesions are not evenly distributed across networks, but rather preferentially affect the frontotemporal nodes of the limbic network. This lesion correction method can facilitate inclusive, unbiased investigation into the anatomic basis of neurological deficits in patients with TBI and other neuropsychiatric diseases associated with focal lesions.

## Acknowledgments

We thank Cheuk Y. Tang, Ph.D. for assistance with acquisition of MRI data. The LETBI Project is supported by the National Institutes of Health/ National Institute for Neurological Disorders and Stroke and National Institute of Child Health and Development (U01 NS086625 and RF1NS115268). This research was also supported by the NIH Director’s Office (DP2HD101400), National Center for Research Resources (U24RR021382), the National Institute for Biomedical Imaging and Bioengineering (P41EB015896, R01EB006758, R21EB018907, R01EB019956, R01EB023281), the National Institute on Aging (AG022381, R01AG008122, R01AG016495, R01AG008122, U01AG006781, R21AG046657, P41RR014075, P50AG005136), the National Center for Alternative Medicine (RC1 AT005728-01), the National Institute for Neurological Disorders and Stroke (K23NS094538, R21NS109627, R01NS052585, 1R21NS072652, 1R01NS070963, R01NS083534, 5U01NS086625), the Eunice Kennedy Shriver National Institute of Child Health and Human Development (K01HD074651, R01HD071664), and the National Institute on Disability Independent Living and Rehabilitation Research (H133B040033). This research also utilized resources provided by National Institutes of Health shared instrumentation grants S10RR023401, S10RR019307, and S10RR023043. Additional support for this project comes from the James S. McDonnell Foundation, the Nancy and Buster Alvord Endowment, the Rappaport Foundation, the Tiny Blue Dot Foundation, institutional funds from the University of Washington School of Medicine, and the Seton Brain Research Fund.

