## Supplementary Material for "Optimizing the Accuracy of Cortical Volumetric Analysis in Traumatic Brain Injury"

* co-senior authors

### Supplementary Methods and Results

*Classification of Traumatic Brain Injury Severity*

TBI severity was classified according to the Department of Defense classification system[1] based on all available information, including medical records, radiology reports, and self-report as elicited by structured lifetime TBI screening questionnaire (the Brain Injury Screening Questionnaire; BISQ).[2] When duration of unconsciousness was not known and records were not available to confirm presence of intracranial abnormality, severity was coded as missing.

*Lesion Correction – Methodological Principles*

This lesion correction method differs in several ways from the standard FreeSurfer process of editing the brain mask and applying control points. Generating cortical volume measurements requires FreeSurfer to model two surfaces, the grey matter (GM) surface and the white matter (WM) surface. Each surface mesh is comprised of thousands of vertices, with 1:1 pairings of vertices between the two surfaces. The distance between these two surfaces at any given vertex-pair provides a measure of cortical thickness. To compute the volume of a cortical region (such as a network of the 7-Network Yeo atlas), FreeSurfer computes the average regional thickness and then multiplies that value by the region's surface area. Thus, any defect or inaccuracy in these surfaces will yield inaccurate volumetric measures.

Historically, standard editing of the FreeSurfer-generated GM and WM surfaces has involved the manual creation of "control points" (typically by a trained research technician). These control points are used to modify the surface models, thereby improving the anatomic accuracy of the surface meshes (for both GM and WM). However, there are fundamental limitations to this approach. Most importantly, this approach relies upon the assumption that the brain has normal anatomy and that any surface inaccuracies are related to the FreeSurfer processing pipeline. Yet, in patients with cortical lesions, such as the patients studied here, the lesions create surface inaccuracies that are attributable to anatomic distortions and encephalomalacia, independent of inaccuracies related to the FreeSurfer processing pipeline. When these cortical lesions are present, the GM and WM surfaces are often undetectable on T1-weighted images (as shown in Figure 1), even when using a high-resolution MEMPRAGE sequence. Hence, any FreeSurfer-generated measurement of the cortical thickness is baseless. Adding control points can move the surface models closer to a rater's desired location, but it is unlikely that the resulting cortical volume measures would be anatomically accurate or biologically valid. Furthermore, nearly all aspects of the FreeSurfer software package depend on a continuous surface mesh, thereby assuming a complete and undamaged cortex. In patients with lesions caused by TBI, there may be encephalomalacic regions of cortex that disrupt cortical continuity. In short, when cortical lesions alter the GM and WM surfaces, the control-point editing method does not generate accurate cortical volume measurements.

This methodological limitation of the control-point approach is the main motivation for the lesion correction method proposed here. The proposed method makes no assumptions about the GM and WM surface anatomy at sites of cortical lesions, and it minimizes bias by requiring the manual rater simply to identify inaccurate surfaces without changing the surfaces in a subjective manner. The method maintains the continuity of the reconstructed FreeSurfer surface mesh while also accounting for regions of cortical atrophy.

*SNR/CNR Correlation with Qualitative Assessments of Scan Quality*

Due to the presence of severe anatomic distortions, two of the 98 patients' scans did not complete FreeSurfer's standard processing pipeline (visual assessment scores=0). To explore the relationship between visual and quantitative assessments of scan quality, we measured the signal-to-noise ratio (SNR) and contrast-to-noise ratio (CNR) in the remaining 96 scans that completed the FreeSurfer reconstruction process. We calculated SNR using the white-matter (WM) segmentation, and we calculated CNR using the WM-GM and GM-cerebrospinal fluid contrasts (see https://github.com/ComaRecoveryLab/Lesion_Correction for specific commands). We tested the hypothesis that the quantitative SNR and CNR scores differed between the groups of scans that received visual assessment scores of 1, 2, and 3.

Of the 96 scans, nine received a visual assessment score of 1 by the two raters and were excluded, yielding a final sample size of 87 patients. All 87 patients were assigned a visual quality score of 2, indicating the need for minor FreeSurfer editing (e.g. editing of the brain mask and use of control points). A two-tailed T-test demonstrated that the quantitative SNR but not the CNR values differed between the 87 patients included in the analysis and the 9 patients excluded ($p<0.0001$ and $p=0.65$, respectively). A summary of the quantitative scores for the scans that were excluded versus the scans that were included is provided in Supplementary Table 2.

*Lesion Correction Protocol and Guide*

The newly proposed lesion correction method involves the following steps, which can be performed by a research technician in less than 60 minutes of active time per patient. The example we have provided can be applied to any surface parcellation already registered to a subjects' FreeSurfer surfaces. If the research technician has already run recon-all and performed the classic manual edits, skip to step (3). All code relating to the steps described below is distributed at https://github.com/ComaRecoveryLab/Lesion_Correction.

1) Process the patient's MRI through FreeSurfer's standard recon-all pipeline. If applicable, use the "-bigventricles" option to improve anatomical segmentation.

2) Spend approximately 30 minutes assessing FreeSurfer output and applying manual edits where necessary before re-running recon-all to apply the adjustments. Repeat this step as needed. For additional details on troubleshooting a FreeSurfer output, see the FreeSurfer Guide to Recon Editing https://surfer.nmr.mgh.harvard.edu/fswiki/FsTutorial/TroubleshootingData.

3) Using the Development Version of FreeView (free download available https://surfer.nmr.mgh.harvard.edu/pub/dist/freesurfer/dev), load the patient's T1.mgz and pial surface files. To avoid unnecessary distractions, turn the curvature off and hide 3D slices. A template terminal command is provided here:

```
freeview \
-v $SUBJECTS_DIR/<Subject ID>/mri/T1.mgz \
-f $SUBJECTS_DIR/<Subject ID>/surf/lh.pial:curvature_method=off \
-f $SUBJECTS_DIR/<Subject ID>/surf/rh.pial:curvature_method=off \
-hide-3d-slices
```

4) Select the "1 & 3 Horizontal" layout from the view panel selection on the top toolbar. This can also be called in the command displayed above by including "-layout 4" at the end.

5) Scroll through the T1.mgz to identify any reconstructed surfaces that pass through cortical lesions (see the Fig. 1 for example). Make note of these lesions, as you will manually label each one in the following steps.

6) Left-click on the "Custom Fill" button located under the loaded surface files on the left-hand toolbar. Then, select "Make Path" and left-click the 3D surface render to place points outlining the lesion on the ??.pial surface (where "??" stands for "rh" or "lh"

depending on the hemisphere). It is very important that these points should maintain an unbroken chain and only be placed on a single hemisphere.

7) Once the point-outline is complete, left-click on the “Make Closed Path” button to connect the points.

8) Now that a lesion path has been created, remove the points by left-clicking “Clear Marks”. Left-click to create a single point anywhere inside the closed path and then left-click “Custom Fill” (followed by “Fill”, when prompted) to fill the lesion label (Fig. 1, middle row).

9) The label will now be filled and a new label file (“label_?”) will appear in the Label Index on the left-hand toolbar. Left-click “Save” and assign the label a new name with the prefix “lh.” or “rh.” to specify the hemisphere, followed by “lesion-??.label” (where “??” denotes a 0-padded integer between 00 and 99) to note the number of previous labels on that hemisphere. For example, the first lesion labeled on the left hemisphere of any given subject will be named, “lh.lesion-01.label”. Repeat this step for every cortical lesion that has interfered with either of the pial surfaces.

10) The “correct_segstats.sh” wrapper script provided on GitHub should automatically combine your labels for each subject’s hemisphere into overlays (“??.all-lesions.mgz”) used to correct the volumetric statistics calculated by FreeSurfer. In short, the script combines each subject’s label files into a single annotation file for each hemisphere and then converts the annotation files into overlay files using the “mris_label2annot” and “mris_annotation2label” commands, respectively. Then, the script applies the “mris_anatomical_stats” command to extract the cortical volume measurements and ignore any anatomical region within the lesion overlay file. The script will then curate

each subjects stats file into two stats tables (one for each hemisphere) by implementing the “aparcstats2table” command.

**Supplementary Table 1. Patient Clinical and Demographic Characteristics**

| | Patients with Cortical Lesions (n=46) | Patients without Cortical Lesions (n=41) | Patients with Unanalyzable MRI (n=11) | Patients Excluded from MRI (n=42) |
|---|---|---|---|---|
| **Age (mean +/- SD years)** | 58.5 +/- 11.2 | 54.8 +/- 12.8 | 61.3 +/- 10.4 | 57.1 +/- 14.8 |
| **Sex (M/F)** | 26/20 | 27/14 | 8/3 | 26/16 |
| **Years from Most Recent TBI to MRI (mean +/- SD)** | 10.1 +/- 11.2 | 11.9 +/- 10 | 16.5 +/- 16.1 | Not applicable |

**Supplementary Table 2. Qualitative and Quantitative Assessment of MRI Data**

| | Inclusion Cohort (n=87) | Exclusion Cohort (n=9)* | P Value |
|---|---|---|---|
| **SNR** | 15.19 +/- 3.35 | 8.26 +/- 2.74 | <0.0001 |
| **CNR** | 0.99 +/- 0.31 | 0.94 +/- 0.18 | 0.65 |

* For two subjects, the FreeSurfer processing pipeline could not be completed. Thus, only 9 of the 11 excluded patients could undergo SNR and CNR analysis.
P values were determined using two-tailed T-tests.

### Supplementary Table 3. Network Overlap with Cortical Lesions

| Network | Number of Lesions Within Network (n=374)* | Number of Patients with Lesioned Network (of 46 Patients with Cortical Lesions) | Average % of Network Surface Area Affected by Overlapping Lesions |
|---|---|---|---|
| **Default Mode** | 93 | 44 (95.7%) | 1.4 +/- 2.1 |
| **Salience** | 60 | 33 (71.7%) | 1.2 +/- 2.0 |
| **Limbic** | 89 | 44 (95.7%) | 4.4 +/- 3.7 |
| **Dorsal Attention** | 20 | 16 (34.8%) | 0.8 +/- 1.2 |
| **Executive Control** | 58 | 36 (78.3%) | 1.6 +/- 2.4 |
| **Somatomotor** | 45 | 31 (67.4%) | 1.3 +/- 2.1 |
| **Visual** | 9 | 9 (19.6%) | 1.3 +/- 1.7 |

* The total number of lesions within all networks is greater than 120 because most lesions overlapped with more than one network. On average, lesions overlap with mean +/- SD 4.6 +/- 1.6 of the 7 networks.

% lesion overlap data are reported as mean +/- SD

**Supplementary Table 4. Network-specific Effects of Cortical Lesions on Average Cortical Volumetric Measures**

| Network | Pre-Correction Cortical Volume (ml) | Post-Correction Cortical Volume (ml) | % Change in Cortical Volume* |
|---|---|---|---|
| **Default Mode (n=44)** | 67.4 +/- 25.8 | 70.6 +/- 26.2 | 5.3 +/- 6.1 |
| **Salience (n=33)** | 30.8 +/- 12.6 | 31.5 +/- 12.4 | 3.8 +/- 4.7 |
| **Limbic (n=44)** | 26.4 +/- 11.5 | 30.1 +/- 13.0 | 12.7 +/- 9.7 |
| **Dorsal Attention (n=16)** | 31.7 +/- 12.1 | 31.9 +/- 12.0 | 2.2 +/- 3.0 |
| **Executive Control (n=36)** | 39.5 +/- 15.8 | 40.8 +/- 16.0 | 4.5 +/- 5.5 |
| **Somatomotor (n=31)** | 44.6 +/- 17.3 | 45.6 +/- 17.1 | 3.7 +/- 5.4 |
| **Visual (n=9)** | 49.5 +/- 18.6 | 49.7 +/- 18.7 | 1.8 +/- 2.1 |

This table summarizes the percent volume change in each network caused by the lesion correction method. The average volume (ml) for each network of the 7-Network Yeo Atlas is provided for the 46 patients with cortical lesions. The limbic network was disproportionately lesioned compared to other networks due to the frontotemporal distribution of cortical lesions, as visualized in Figure 2.

* Measurements for the percent change in cortical volume include networks that have been impacted by the volume correction method. See column 3 of Supplementary Table 3 for the number of paired (pre vs. post) volumes that were used to calculate each network's percent change in cortical volume.

### Video Legends

#### Video 1. Overview of Network-based Lesion Overlap Analysis

A left temporal lesion is shown in yellow for a representative patient. The yellow lesion mask represents all of the cortical vertices that were affected by the cortical lesion, as evidenced by inaccuracies in the FreeSurfer-generated cortical surface. Next, the 7-Network Yeo atlas is shown on the cortical surface, with the same network color-labeling shown in Figure 2. Finally, the portions of the Yeo networks that overlap the lesion mask are shown. This video demonstrates how a single lesion can overlap with multiple functional networks.

#### Video 2. A Three-Dimensional View of Lesion Topology on the Cortical Surface

The lesion topology map shown in the left panel of Figure 2 is displayed here in three dimensions. The anatomic regions most commonly affected by cortical lesions were the frontal and temporal lobes, particularly the frontal poles, temporal poles and orbitofrontal regions.

#### Video 3. Visualization of Individual Lesion Topology on the Cortical Surface

Motivated by prior efforts to map the cortical topology of traumatic contusions by Courville in 1942,[3] we show the sequential overlapping of every cortical lesion identified in the 46 patients with lesions in this study. The video is designed in the style of Courville's classic lesion topology figure.

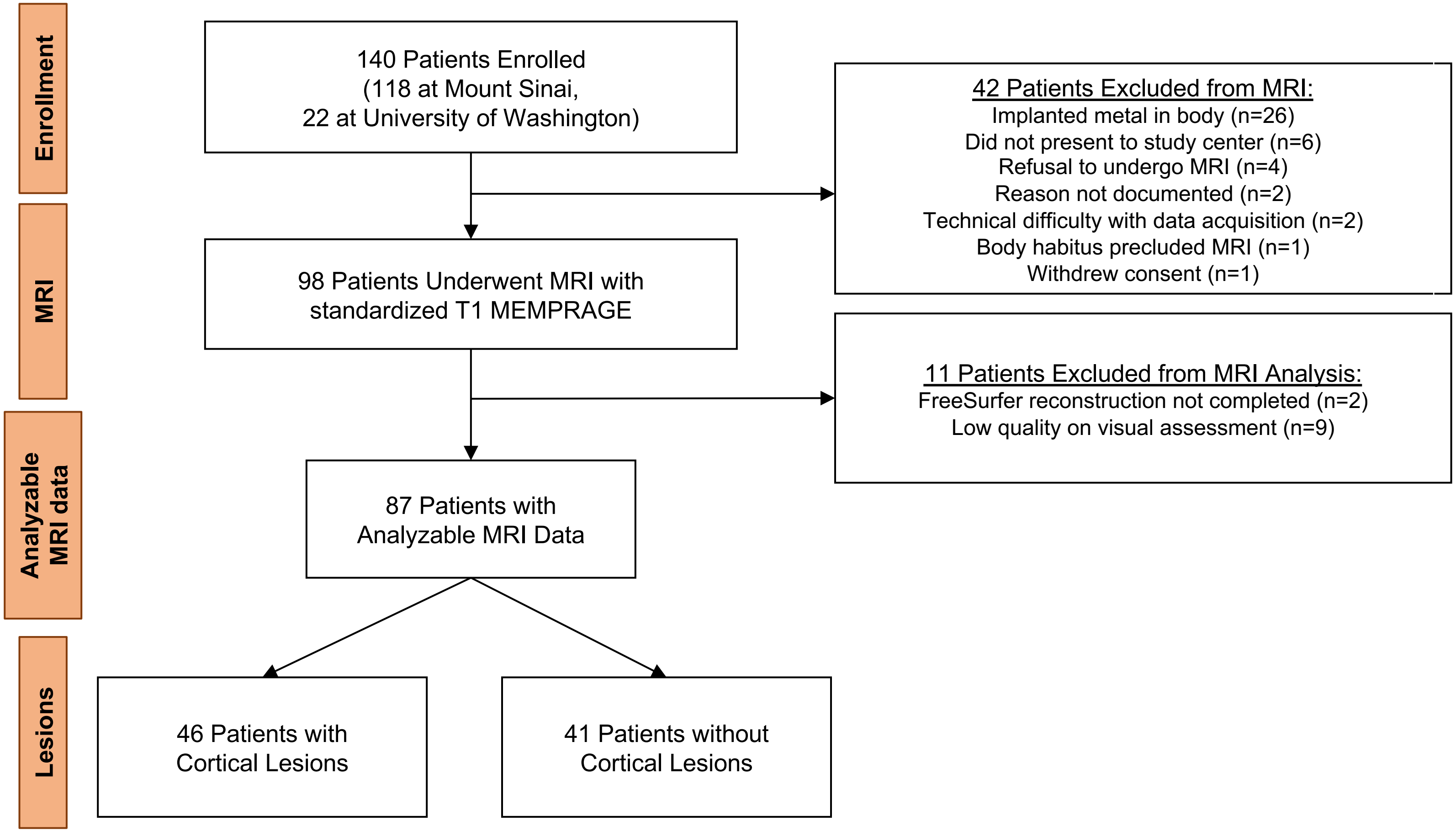


**Supplementary Figure 1. CONSORT Diagram.**

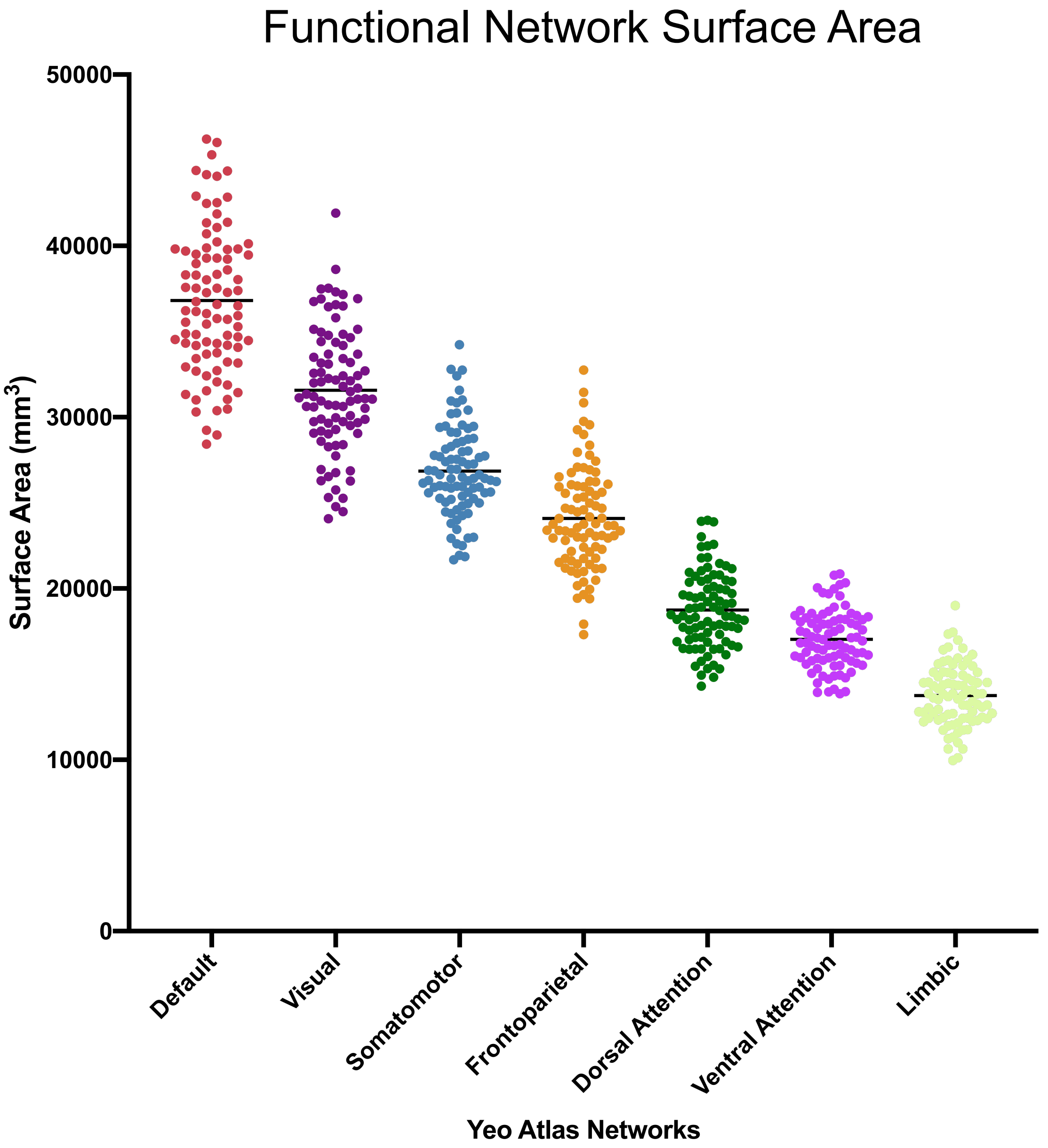


**Supplementary Figure 2. Average Surface Area Measures for the 7 Yeo Functional Networks Across all Patients.** The limbic network has the smallest average surface area of the 7 networks, yet is the network that is most commonly affected by cortical lesions in patients with chronic moderate-to-severe traumatic brain injury.
